# Heh2/Man1 may be an evolutionarily conserved sensor of NPC assembly state

**DOI:** 10.1101/2020.06.29.178129

**Authors:** Sapan Borah, David J. Thaller, Zhanna Hakhverdyan, Elisa C. Rodriguez, Michael P. Rout, Megan C. King, C. Patrick Lusk

**Affiliations:** Department of Cell Biology, Yale School of Medicine, 295 Congress Avenue, New Haven, CT, 06520; The Rockefeller University, 1230 York Avenue, New York, NY, 10065

## Abstract

Integral membrane proteins of the Lap2-emerin-MAN1 (LEM) family have emerged as important components of the inner nuclear membrane (INM) required for the functional and physical integrity of the nuclear envelope. However, like many INM proteins, there is limited understanding of the biochemical interaction networks that enable LEM protein function. Here, we show that Heh2/Man1 can be affinity purified with major scaffold components of the nuclear pore complex (NPC), specifically the inner ring complex, in evolutionarily distant yeasts. Interactions between Heh2 and nucleoporins is mediated by its C-terminal winged-helix (WH) domain and are distinct from interactions required for INM targeting. Disrupting interactions between Heh2 and the NPC leads to NPC clustering. Interestingly, Heh2’s association with NPCs can also be broken by knocking out Nup133, a component of the outer ring that does not physically interact with Heh2. Thus, Heh2’s association with NPCs depends on the structural integrity of both major NPC scaffold complexes. We propose a model in which Heh2 acts as a sensor of NPC assembly state, which may be important for NPC quality control mechanisms and the segregation of NPCs during cell division.

## Introduction

The eukaryotic genome is enclosed by a nuclear envelope that is contiguous with the endoplasmic reticulum (ER). Despite this continuity, the nuclear envelope contains a unique proteome that defines its function as a selective barrier. This barrier not only establishes nuclear-cytoplasmic compartmentalization but also directly impacts genome organization and function at the nuclear periphery (Mekhail and Moazed, 2010; Taddei and Gasser, 2012; Buchwalter et al., 2019). The key elements of this biochemical specialization are the nuclear pore complexes (NPCs), which control nucleocytoplasmic molecular exchange, and proteins specifically associated with the inner and outer nuclear membranes (INM and ONM)(Ungricht and Kutay, 2017; Hampoelz et al., 2019). While ONM proteins generally act as adaptors that connect the cytoskeleton to the nucleus (Burke and Roux, 2009), INM protein function is less well defined. This is due in part to challenges inherent with defining biochemical interactions between low abundance integral membrane proteins that exist within a complex and integrated network of peripheral chromatin and nuclear scaffold proteins like the lamins (outside of yeasts). Nonetheless, there is confidence that there are several dozen integral INM proteins with the most evolutionarily conserved families being the LAP2-emerin-MAN1 (LEM) proteins and the SUN family proteins (Mans et al., 2004; Ungricht and Kutay, 2015).

LEM family proteins are so named for their LEM domain, a short ~40 amino acid helix-extension-helix motif that, at least in higher eukaryotes, binds to barrier to autointegration factor (BAF)(Furukawa, 1999; Cai et al., 2007). As there is no BAF in yeasts, the LEM domain must possess other conserved functions, which may more directly relate to genome integrity, ensuring the stability of repetitive DNA (Mekhail et al., 2008), and also contributing to the mechanical integrity of the nucleus (Schreiner et al., 2015). There are up to seven LEM domain proteins in humans but in the two most commonly used yeast models, *Saccharomyces cerevisiae* (Sc) and *Schizosaccharomyces pombe* (Sp) there are only two: ScHeh1(Src1)/SpHeh1(Lem2) and ScHeh2/SpHeh2(Man1)(Barton et al., 2015). Of these two, ScHeh1 and SpHeh1 are likely orthologs derived from a common ancestor, while ScHeh2 and SpHeh2 resulted from independent duplication events of their respective paralogs ScHeh1 and SpHeh1 (Rhind et al., 2011; Gonzalez et al., 2012). Despite their independent evolutionary history, there is evidence that Heh2 in both yeasts specifically makes functional connections with NPCs. For example, in *S. cerevisiae*, we demonstrated synthetic genetic interactions between genes encoding NPC components (nucleoporins or nups), and *HEH2* (Yewdell et al., 2011). In the *S. pombe* cousin, *S. japonicus*, it has also been suggested that Heh2 supports connections between chromatin and NPCs to support their segregation between daughter cells in mitosis (Yam et al., 2013). However, the underlying biochemical connections between Heh2 and the NPC are not understood.

Understanding the nature of the connections between Heh2 and the NPC may also help illuminate mechanisms underlying the biogenesis of NPCs. As the total proteome, interactome and structure of NPCs have come to light, it is now understood that the enormous (50-100 MD) NPC is built from a relatively small (~30) number of nups (Hampoelz et al., 2019). These nups are organized into modular subcomplexes that, in multiples of 8, assemble the 8-fold radially symmetric NPC scaffold composed of inner and outer ring complexes (IRC and ORC), the central transport channel and asymmetric (perpendicular to the plane of the nuclear envelope) cytosolic filaments/mRNA export platform and nuclear basket (Kosinski et al., 2016; Kim et al., 2018). How NPCs are assembled in space and time during interphase remains ill-defined, but likely begins within the nucleus at the INM (Marelli et al., 2001; Makio et al., 2009; Yewdell et al., 2011; Mészáros et al., 2015; Otsuka et al., 2016). The recruitment of nups to an assembly site occurs alongside membrane-remodeling that evaginates the INM and ultimately drives fusion with the ONM (Otsuka et al., 2016). Consistent with an inside-out model, the cytosolic-facing mRNA export platform is likely added at a terminal step in NPC assembly (Otsuka et al., 2016; Onischenko et al., 2017). In genetic backgrounds where the cytoplasmic-facing mRNA export platform is not assembled, herniations or blebs are observed over assembling NPCs, which may reflect defects in INM-ONM fusion and/or the triggering of NPC assembly quality control pathways (Thaller and Lusk, 2018).

Both Heh1 and Heh2 have been implicated in mechanisms of NPC assembly quality control in which they regulate the recruitment of the endosomal sorting complexes required for transport (ESCRT) to the nuclear envelope (Webster et al., 2014, 2016; Thaller et al., 2019). One early model suggested that Heh2 may differentially bind to NPC assembly intermediates over fully formed NPCs (Webster et al., 2014). However, this has yet to be formally interrogated. In order to be more incisive as to how Heh2 impacts NPC function, here we have thoroughly analyzed the biochemical interaction network of endogenous Heh2. Using two evolutionary distant yeasts, we show that Heh2 can co-purify with the NPC’s IRC. These interactions do not require the LEM domain or any INM targeting sequences but instead depend on a C-terminal domain predicted to fold into a winged helix (WH)(Caputo et al., 2006). Further, by decoupling NPC clustering from perturbations to NPC structure, we demonstrate that Heh2 associates with NPCs *in vivo*. Most interestingly, the association of Heh2 with the NPCs can be completely broken by knocking out Nup133, a nucleoporin of the ORC, suggesting that Heh2’s association with the NPC depends on its structural integrity. Taken together, we suggest a model in which Heh2 may be a sensor of NPC assembly state.

## Results

### Heh2 binds to specific nups in evolutionarily distant yeasts

To better define the interacting partners of Heh1 and Heh2, we performed one-step affinity purifications of Heh1-TAP and Heh2-TAP (produced at endogenous levels) from cryolysates derived from logarithmically growing budding yeast (Hakhverdyan et al., 2015). As shown in Fig. 1A, we did not detect any obvious stoichiometric binding partners of Heh1-TAP despite robust recovery of the fusion protein. In marked contrast, Heh2-TAP co-purified with at least 8 additional proteins, which were visible by SDS-PAGE and Coommassie blue staining of bound fractions. Excision of these bands followed by mass spectrometric (MS) protein identification revealed that Heh2 binds to the IRC of the NPC and a subset of cytosolic-facing nups, including Nup159, Nup188, Nup192, Nup170, Pom152, Nup157, Nup116, Nic96 and Nsp1. For context, we have colored the identified nups in a diagram of a single spoke from the budding yeast NPC structure (Kim et al., 2018) in Fig. 1A.

**Figure1.**
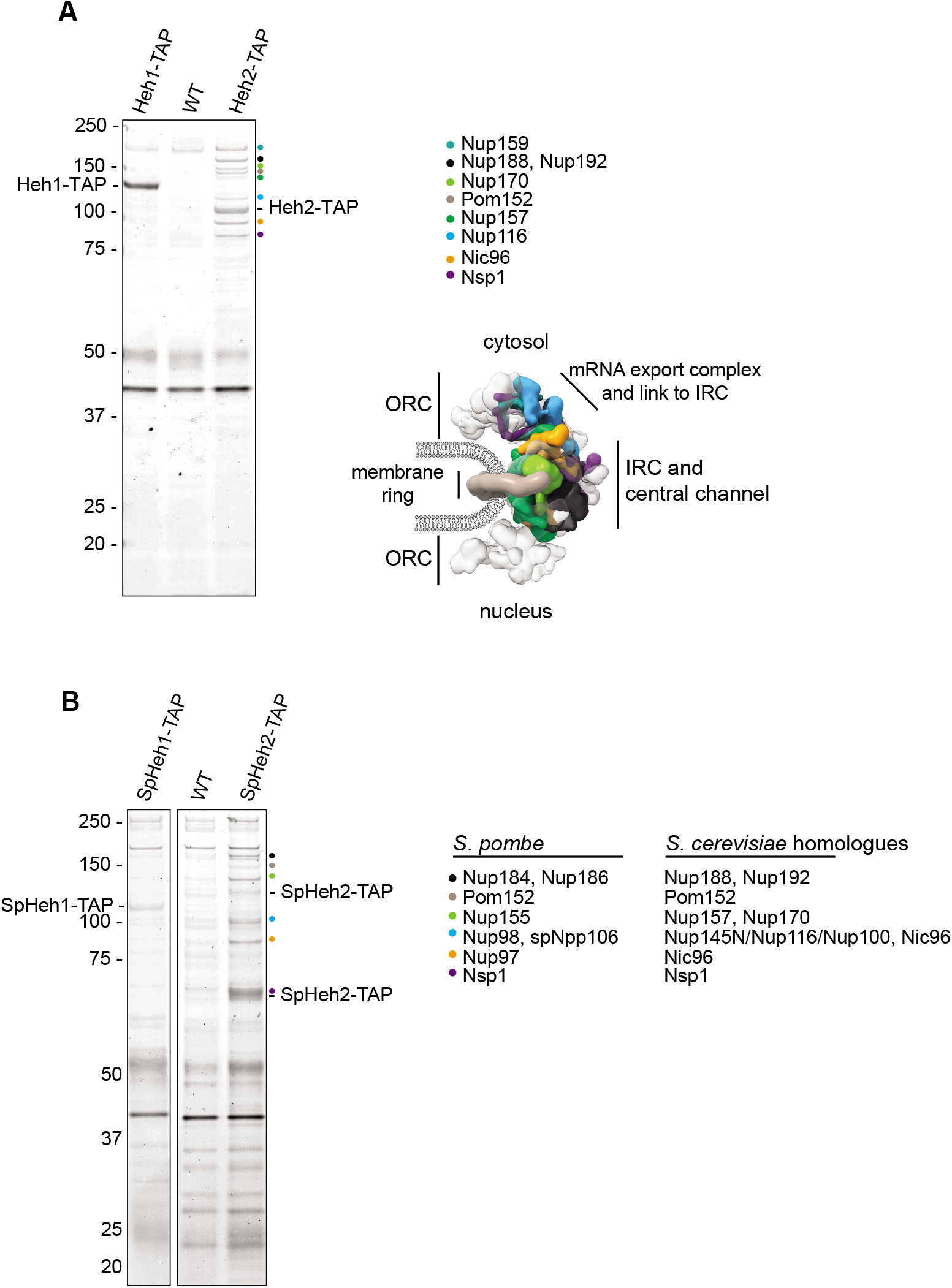
Heh2 binds to specific nups in evolutionary distant yeasts. **(A)** Heh2 specifically binds the IRC. Affinity purifications were performed from cell extracts derived from strains expressing endogenous Heh1-TAP or Heh2-TAP or from WT cells (no TAP). Bound proteins were separated by SDS-PAGE and visualized by Coomassie staining. Numbers at left indicate position of MW standards in kD. Heh1-TAP and Heh2-TAP are indicated, and colored circles demark proteins identified by MS from Heh2-TAP lane, as indicated in key. This color scheme is also used to indicate positions of nups within a single spoke of the NPC structure (from PDBDEV_00000010; Kim et al., 2018). ORC is outer ring complex, IRC is inner ring complex. **(B)** As in A but affinity purifications performed from *S. pombe* cell extracts. The corresponding *S. cerevisiae* homologues of the identified *S. pombe* nups are also listed.

We were next curious whether Heh2’s association with the NPC was also observed in other yeast species where the NPC structure is different than in budding yeast. For example, fission yeast NPCs are made up of a similar catalogue of nups (Baï et al., 2004; Chen et al., 2004; Asakawa et al., 2014), but there is evidence that there is asymmetry with respect to the ORC, which contains 16 copies (instead of 8) of the “Y” complex on the nucleoplasmic side of the NPC (Asakawa et al., 2019). Of additional interest, although *HEH1* in both *S. cerevisiae* and *S. pombe* is derived from a common ancestor, these yeasts are separated by ~500 million years of evolution (Rhind et al., 2011). Intriguingly, and in contrast, *ScHEH2* and *SpHEH2* arose from distinct duplication events (Mans et al., 2004), and might therefore be expected to carry out distinct functions.

Interestingly however, despite this unique evolutionary history, the affinity-purifications of SpHeh2-TAP and SpHeh1-TAP were qualitatively similar to the *S. cerevisiae* versions with SpHeh1-TAP co-purifying with few specific proteins (compare to the WT control) and SpHeh2-TAP with several specific species (Fig. 1B). Note that SpHeh2-TAP is proteolytically sensitive and is purified both as a full length (~115 kDa) and a smaller (~65 kDa) form (Fig. 1B). Nonetheless, like its distant *S. cerevisiae* cousin, the SpHeh2-complex consisted of essentially the same subset of inner ring nups including Nup184, Nup186, Nup155, Pom152, Npp106, Nup98 and Nup97 (Fig. 1B). To facilitate a comparison, the *S. cerevisiae* homologues are listed next to the identified *S. pombe* nups in Fig. 1B. Thus, despite the distinct duplication events that gave rise to *HEH2* in both species, the physical association of Heh2 with the IRC likely points to an important and conserved function that was likely shared by a common ancestor before being independently specialized in the two species lineages.

### Heh2 fails to interact with NPCs lacking Nup133

That Heh2 binds to nups suggests that it may be a component of the NPC. To assess this possibility, we next examined the distribution of Heh2-GFP at the nuclear envelope alongside an NPC marker, Nup82-mCherry. We also took advantage of a standard approach of knocking out *NUP133*, which leads to NPC clustering and facilitates co-localization analysis, as individual NPCs cannot be resolved with conventional light microscopy (Doye et al., 1994; Pemberton et al., 1995; Li et al., 1995; Aitchison et al., 1995; Heath et al., 1995). Consistent with prior work (Yewdell et al., 2011), we observed a punctate NPC-like distribution of Heh2-GFP at the nuclear envelope of otherwise WT cells, which exhibited some co-localization with Nup82-mCherry (Fig. 2A). Indeed, when we quantified the correlation between the GFP and mCherry fluorescence at each pixel along the nuclear envelope of 20 cells, we observed a modest positive correlation (*r* = 0.39; Fig. 2B). In marked contrast, deletion of *NUP133* led to a striking anti-correlation between Nup82-mCherry and Heh2-GFP (*r* = - 0.27), which was obvious in the micrographs where Heh2-GFP was diminished or undetectable at the Nup82-mCherry clusters (Fig. 2A, B, bottom panels). We note further that Heh2-GFP is no longer punctate along the nuclear envelope in *nup133*Δ cells, which suggests that there may in fact be an association with NPCs (as supported by the biochemistry) but that this interaction is broken without Nup133.

**Figure 2.**
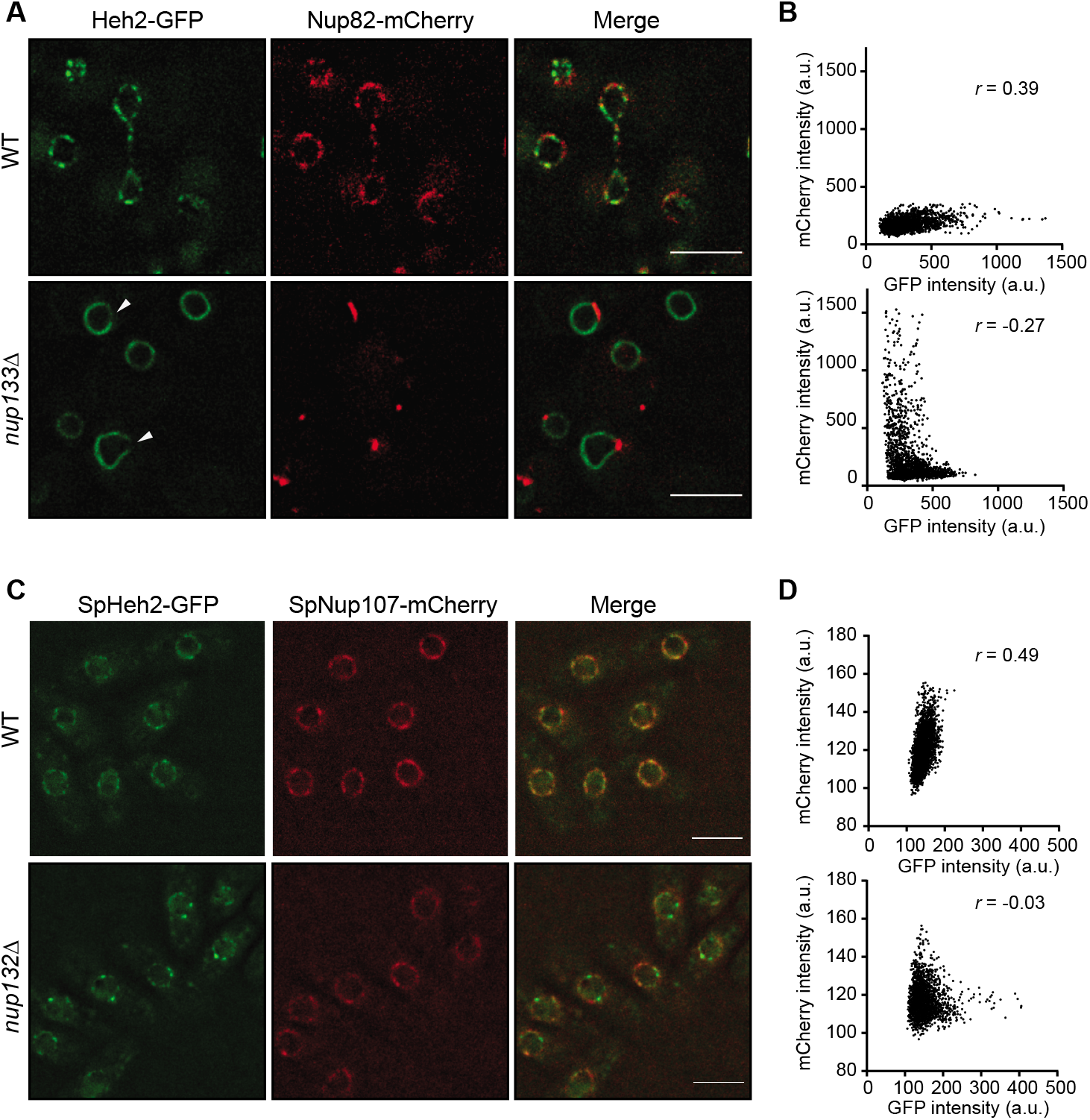
Heh2 fails to interact with NPCs lacking Nup133. **(A)** Deconvolved fluorescence micrographs of Heh2-GFP and Nup82-mCherry with merge in WT and *nup133*Δ strains. Arrowheads point to regions depleted of Heh2-GFP that contain Nup82-mCherry in a cluster. Scale bar is 5 μm. **(B)** Scatterplot with Pearson correlation coefficient (*r*) of Heh2-GFP and Nup82-mCherry fluorescence intensity (in arbitrary units, a.u.) along the nuclear rim of 20 cells, from two independent experiments. **(C)** Deconvolved fluorescence micrographs of SpHeh2-GFP, and SpNup107-mCherry with merge in WT and *nup132Δ S. pombe* cells. Scale bar is 5 μm. **(D)** Scatterplot with Pearson correlation coefficient (*r*) of SpHeh2-GFP and SpNup107-mCherry fluorescence intensity (in arbitrary units, a.u.) along the nuclear rim of 20 cells, from two independent experiments.

To continue with the exploration of potential functional commonalities between ScHeh2 and SpHeh2, we also tested whether deletion of the orthologous *S. pombe Nup132* impacted SpHeh2-GFP distribution (Baï et al., 2004). As has been reported by others, SpHeh2 also has a punctate distribution evocative of NPCs (Fig. 2C)(Gonzalez et al., 2012; Steglich et al., 2012). Consistent with this, we observed coincidence between SpHeh2-GFP and SpNup107-mCherry fluorescence with a correlation value of *r* = 0.49 (Fig. 2D, top). Interestingly, as in *S. cerevisiae*, deletion of *Nup132* lead to a clear anti-correlation (*r* = - 0.03) of the SpHeh2-GFP and SpNup107-mCherry signals, suggesting that their physical interaction could be disrupted (Fig. 2D, bottom). Remarkably, this anti-correlation was observed even with minimal clustering of SpNup107-mCherry in this strain (Fig. 2C). Thus, this result reinforces that disrupting NPC structure by deleting a critical ORC component compromises Heh2’s ability to interact with NPCs in both organisms.

### Heh2 co-localizes with NPCs

To reconcile the apparent inconsistency between the affinity purifications, which suggested that Heh2 binds NPCs, and the lack of Heh2-GFP co-clustering with nups in *nup133*Δ strains, we sought an orthogonal approach to assess Heh2-GFP co-localization with NPCs that were not missing key structural components. In prior work, we observed that the anchor-away approach (Haruki et al., 2008)(Fig. 3A) can drive rapid NPC clustering through the rapamycin-induced dimerization of a Nsp1-FRB fusion that was incorporated into NPCs (and likely exposed to the cytosol) with Pma1-FKBP12 (a plasma membrane anchor, Fig. 3A) within 15 min (Colombi et al., 2013). The rapidity of this response strongly suggested that fully formed NPCs are driven into clusters independent of NPC mis-assembly. Further, we did not detect any removal of Nsp1-FRB from NPCs under these conditions (Colombi et al., 2013). Consistent with this, we assessed the co-localization of Nup82-GFP with Nup170-mCherry in strains expressing Nsp1-FRB and Pma1-FKPB12 in the presence of carrier alone (DMSO) or rapamycin. As expected, both of the fluorescent proteins localized in a punctate distribution at the nuclear envelope in the presence of DMSO with a significant *r* = 0.48 positive correlation between the GFP and mCherry fluorescence (Fig. 3B, far right panel). Upon addition of rapamycin, we observed rapid clustering and concurrent co-localization of both signals along the nuclear envelope, which was evident in the coincidence of the GFP and mCherry fluorescence peaks of line profiles along the nuclear envelope and a correlation that increased to *r* = 0.74 (Fig. 3B, middle and right panels).

**Figure 3.**
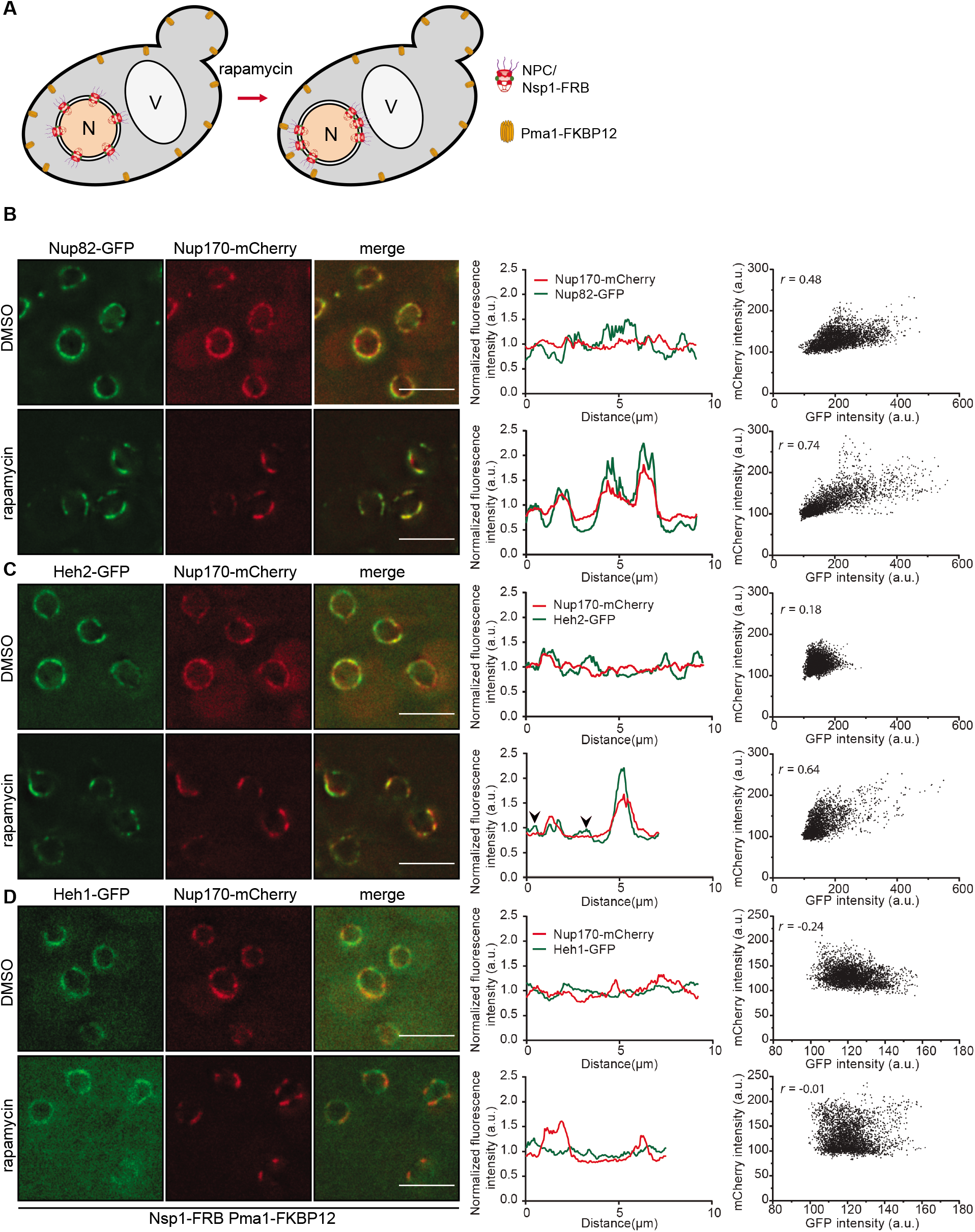
Heh2 associates with NPCs in vivo. **(A)** Schematic of NPC clustering assay mediated by the rapamycin-induced dimerization of Nsp1-FRB (at the NPC) and Pma1-FKBP12. N is nucleus, V is vacuole. **(B-D)** Left: Deconvolved fluorescence micrographs of indicated GFP tagged proteins and Nup170-mCherry as a NPC marker with merge in cells treated with DMSO (carrier) or rapamycin for 15 min. Scale bar is 5 μm. Middle: Line profiles of fluorescence intensity of GFP and mCherry fusions (in arbitrary units, a.u.) along the nuclear envelope of a single cell. Right: Scatterplot with Pearson correlation coefficient (*r*) of GFP and mCherry fluorescence intensity (in arbitrary units, a.u.) along the nuclear rim of 30 cells, from three independent experiments.

We next tested how this approach to NPC clustering influenced Heh2-GFP localization. As a control, we also assessed the distribution of Heh1-GFP, which does not stably interact with nups (Fig. 1A). As shown in Fig. 3C, the addition of rapamycin lead to the clear co-localization of Heh2-GFP and Nup170-mCherry. This again was evident through the examination of line profiles of a representative nuclear envelope where there was coincidence between the peaks of the GFP and mCherry fluorescence and further supported by the increased positive correlation of GFP and mCherry fluorescence (from *r* = 0.18 to *r* = 0.64; Fig. 3C, middle and right panels). Note, however, that unlike the comparison between the two nups (Fig. 3B), there are peaks of Heh2-GFP fluorescence that are not coincident with the NPC clusters (Fig. 3C, arrowheads in line profiles). Thus, while it is clear that Heh2-GFP associates with NPCs, there is also an additional pool of Heh2-GFP at the INM. Last, we did not observe similar effects with Heh1-GFP, which failed to cluster with NPCs (Fig. 3D) or correlate with their distribution (*r* = - 0.01)(Fig. 3D, right panel). Thus, this NPC clustering approach more faithfully mirrored our biochemical analysis of both Heh1 and Heh2 and supports the interpretation that Heh2 is a shared component of NPCs and the INM.

### Inhibition of NPC assembly reduces the Heh2 pool bound to NPCs

A model in which there are two pools of Heh2 was further supported by experiments where we reduced NPC number by inhibiting NPC assembly. For example, by again leveraging the anchor-away strategy, we inhibited NPC assembly by trapping newly synthesized Nup192-FRB-GFP for 3 h (Colombi et al., 2013). Under these conditions, there is a reduction of NPCs that is reflected by lower levels of Nup192-FRB-GFP at the nuclear envelope and a concomitant accumulation of newly synthesized Nup192-FRB-GFP at the plasma membrane (Fig. 4A, B, rapamycin panels). In this scenario, we tested whether Nup192-FRB-GFP and Heh2-mCherry co-localized at the nuclear envelope (Fig. 4B). As a control, we also tested co-localization with Pom152-mCherry (Fig. 4A). While Pom152-mCherry distribution was similar to Nup192-FRB-GFP with line profiles showing coincidence between mCherry and GFP fluorescence peaks along the nuclear envelope (Fig. 4A, far right), there were clear gaps in the Nup192-FRB-GFP signal that were filled by Heh2-mCherry (Fig. 4B, see arrowheads). This result is also represented in line profiles across the nuclear envelope where the Heh2-mCherry signal fills areas that are devoid of GFP-peaks (Fig. 4B, right bottom panel). Importantly, however, a subset of Nup192-FRB-GFP peaks that likely correspond to NPCs that were assembled prior to rapamycin addition still coincided with Heh2-mCherry peaks (Fig. 4B, right bottom panel). Thus, these data are consistent with the interpretation that inhibition of NPC assembly leads to a decrease in the pool of Heh2 bound to NPCs (due to their reduced number) and an increase in the free pool at the INM. This conclusion is further supported by affinity-purifications of Heh2-TAP from Nup192-FRB-GFP strains under the same conditions. While in DMSO-treated conditions the expected IRC profile of nups was detected (Fig. 4C), upon inhibition of NPC assembly with rapamycin, we observed a ~2-3 fold reduction of these nups (orange line in densitometry plot at right) while the total amount of Heh2-TAP affinity purified remained unchanged (Fig. 4C). Thus, we favor a model in which Heh2 remains capable of binding to the IRC in fully formed NPCs, even when their number is decreased upon assembly inhibition.

**Figure 4.**
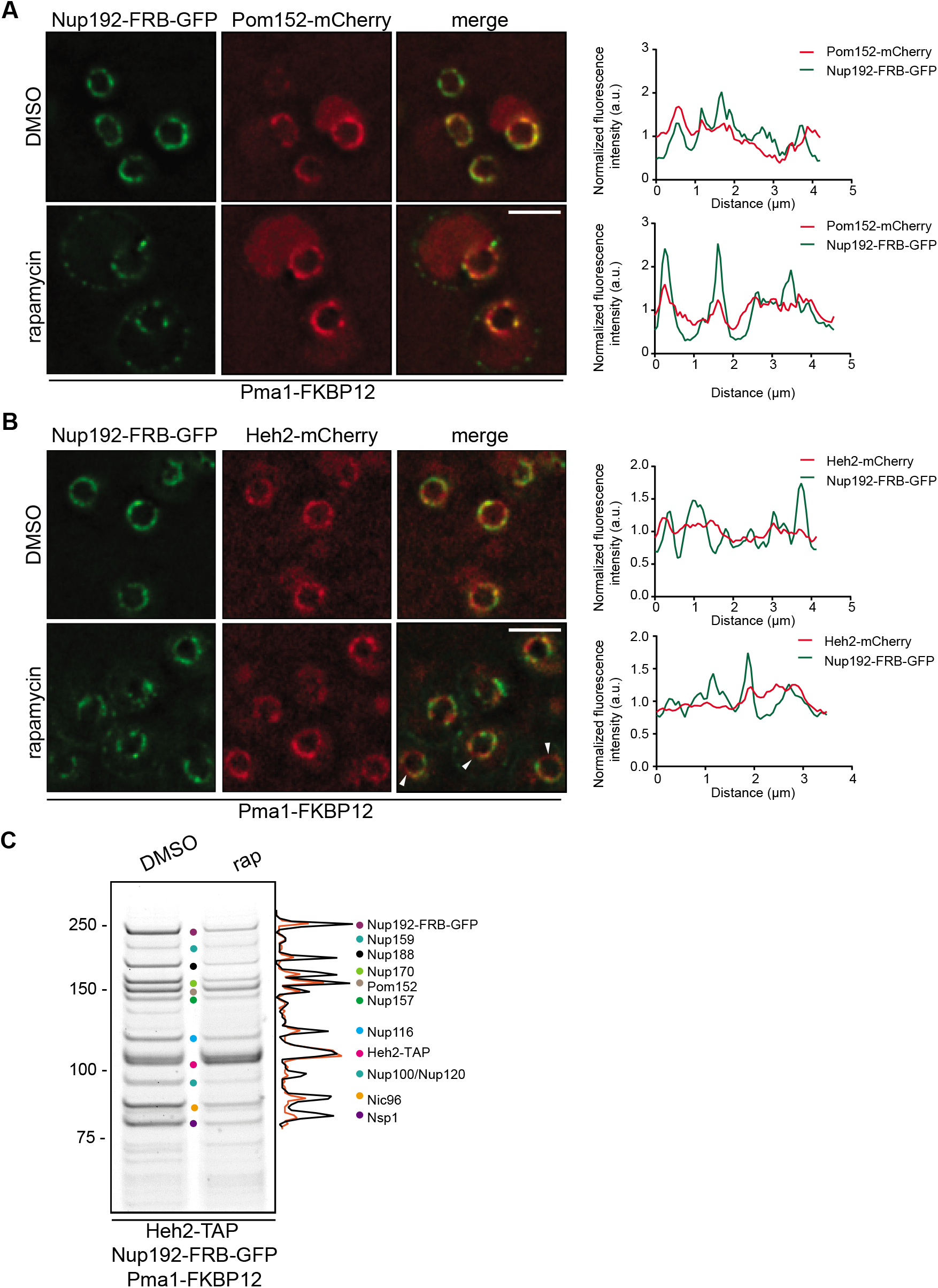
Inhibition of NPC assembly reduces the Heh2-nup bound pool. **(A, B)** Deconvolved fluorescence micrographs of Nup192-FRB-GFP with either Pom152-mCherry or Heh2-mCherry with merge after treating cells with DMSO (carrier) or rapamycin for 3 h to inhibit NPC assembly. Note accumulation of newly synthesized Nup192-FRB-GFP at the plasma membrane as it binds to the Pma1-FKBP12 anchor. Arrowheads point to Heh2-mCherry at the nuclear envelope that is resolvable from Nup192-FRB-GFP signal. Scale bar is 2 μm. At right are line profiles of GFP and mCherry fluorescence intensity (in arbitrary units, a.u.) along the nuclear envelope of single cells corresponding to DMSO (top) and rapamycin (bottom) conditions. **(C)** Inhibiting NPC assembly reduces Heh2-IRC binding. Affinity purifications were performed from cell extracts derived from cells expressing Heh2-TAP with Nup192-FRB-GFP and Pma1-FKBP12 treated with carrier (DMSO) alone, or with rapamycin (rap) to inhibit NPC assembly. Bound proteins were separated by SDS-PAGE and visualized with Coomassie. Position of MW markers (kD) are indicated at left and proteins are marked with colored circles that denote their identity as per key at right. Densitometry of the protein staining of the DMSO (black) and rapamycin (orange) lanes on right.

### Heh2’s association with NPCs depends on the integrity of the NPC scaffold

If Heh2 binds the IRC, it remained unclear why deletion of *NUP133* abrogated Heh2’s NPC association, as the IRC is expected to be intact in this background. Thus, to rule out that Heh2 may be binding IRC nups outside of the context of fully formed NPCs, we directly tested whether deletion of *NUP133* lead to a loss of Heh2 IRC binding. Strikingly, affinity purifications of Heh2-TAP in *nup133*Δ cells did not reveal any obvious binding partners, with the potential exception of Nup159, further supporting the *in vivo* evidence that the structurally deficient *nup133*Δ NPCs are incompetent for binding Heh2 (Fig. 5A). This result is illustrated as a loss of the colored Heh2-interacting nups within the context of a side and center view of a NPC spoke in Fig. 5B. Consistent with the conserved lack of colocalization of scHeh2-GFP and spHeh2-GFP with NPCs in the absence of Nup133/Nup132, we also observed a loss of nups in affinity-purified fractions of SpHeh2-TAP from *nup132*Δ extracts (Fig. 5C).

**Figure 5.**
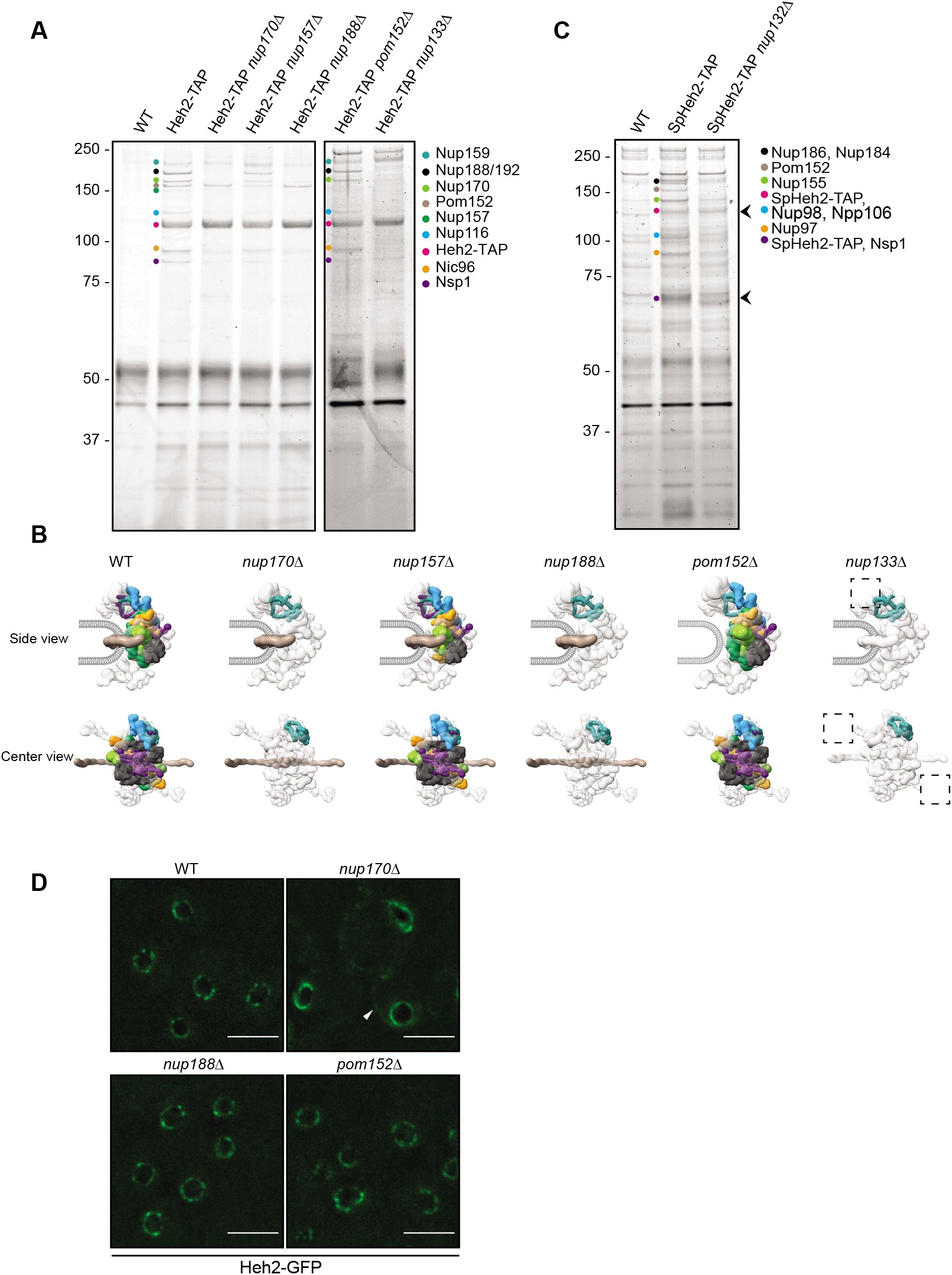
NPC scaffold integrity affects Heh2’s association with NPCs. **(A)** Affinity purifications were performed from cell extracts derived from the indicated nup gene deletion strains expressing endogenous Heh2-TAP or from WT cells (no TAP). Bound proteins were separated by SDS-PAGE and visualized by Coomassie staining. Numbers at left indicate position of MW standards in kD. Proteins are marked with colored circles that denote their identify as per key at right. **(B)** The nups affinity purified from the indicated genetic backgrounds in A are placed within a single spoke of the NPC structure (from PDBDEV_00000010; Kim et al., 2018) in side and center views. Individual nups are colored as in the key in A. **(C)** As in A but affinity purifications performed from *S. pombe* cell extracts. **(D)** Deconvolved fluorescence micrographs of Heh2-GFP in indicated strain backgrounds. White arrowhead points to Heh2-GFP fluorescence at the cortical ER in *nup170*Δ cells. Scale bars are 5 μm.

We next explored the hierarchy of physical interactions that control Heh2’s association with the IRC by affinity-purifying Heh2-TAP from several IRC nup deletion backgrounds. Interestingly, and in contrast with the deletion of *NUP133*, we were unable to define any single knockout of an inner ring nup that fully broke Heh2’s biochemical association with this complex. For example, in cases where we deleted the genes encoding Nup157 or Pom152, we observed the discrete loss of these, and only these, proteins from bound fractions (Fig. 5A, B). Deletion of *NUP170* and *NUP188* led to a more severe disruption of nups bound to Heh2, but in these cases, Pom152 and a band at the molecular weight of Nup159 remained (Fig. 5A, B). Thus, it seems likely that Heh2 makes several direct connections to nups in the IRC, with the most obvious candidates being Pom152, Nup170 and/or Nup188. Heh2 may also directly bind to Nup159, although this association alone is insufficient to maintain association with the NPC *in vivo* (Fig. 2A).

Our inability to fully break interactions between Heh2 and the NPC by abrogating single nups within the IRC was further supported by the lack of any major changes to Heh2-GFP distribution in the *nup170Δ, nup188Δ* and *pom152Δ* strains; in all cases the punctate, NPC-like distribution of Heh2-GFP was retained (Fig. 5D). The one potential exception here was that, in addition to the punctate nuclear envelope distribution, a cortical ER pool of Heh2-GFP could be discerned specifically in *nup170Δ* strains (Fig. 5D, arrowhead). These data are consistent with prior work demonstrating that Nup170 is uniquely required for the efficient targeting of overexpressed Heh2 to the INM (King et al., 2006). Thus, we suggest that, with the exception of Nup170, the physical interactions with the IRC described here are dispensable for INM targeting. Such an assertion is further supported by the exclusive nuclear envelope localization of Heh2-GFP in *nup133*Δ cells where virtually all of its biochemical interactions to the NPC are broken (Fig. 2A). These data thus make the prediction that the INM targeting and NPC-binding elements of Heh2 are distinct.

### The conserved WH domain of Heh2 is required for NPC association

To explore the possibility that INM targeting and NPC-binding may require unique structural elements of Heh2, we generated truncations of Heh2 where the N-terminal nucleoplasmic domain (which contains the INM-targeting information (King et al., 2006; Meinema et al., 2011)) and the C-terminal WH domains are deleted (Fig. 6A). Interestingly, deletion of the N-terminus did not impact binding to nups, as a similar (if more robust) profile of the IRC was recovered in affinity purifications of heh2-(316-663)-TAP (Fig. 6B). These data suggest that Heh2 can reach the NPC (or at least bind to nups) in the absence of its N-terminal INM targeting domain. In marked contrast, deletion of the WH domain, which does not impact INM targeting (Meinema et al., 2011), led to a striking reduction of nup binding (Fig. 6B). These results were also mirrored *in vivo*. For example, compared with Heh2-GFP, heh2-(1-570)-GFP did not exhibit a punctate distribution at the nuclear envelope (Fig. 6C), which was quantified as a reduced coefficient of variation of the fluorescence signal along the nuclear envelope (Fig. 6D). Consistent with the idea that this change in localization of heh2-(1-570)-GFP was due to a loss of its interaction with NPCs, it also failed to cluster with NPCs in the Nsp1-FRB NPC clustering assay (Fig. 6E) with no positive correlation between heh2-(1-570)-GFP and Nup170-mCherry signals in either DMSO (*r* = 0.0) or rapamycin (*r* = - 0.08) treated cells (Fig. 6F). Thus, the WH domain of Heh2 is the major determinant of its association with NPCs.

**Figure 6.**
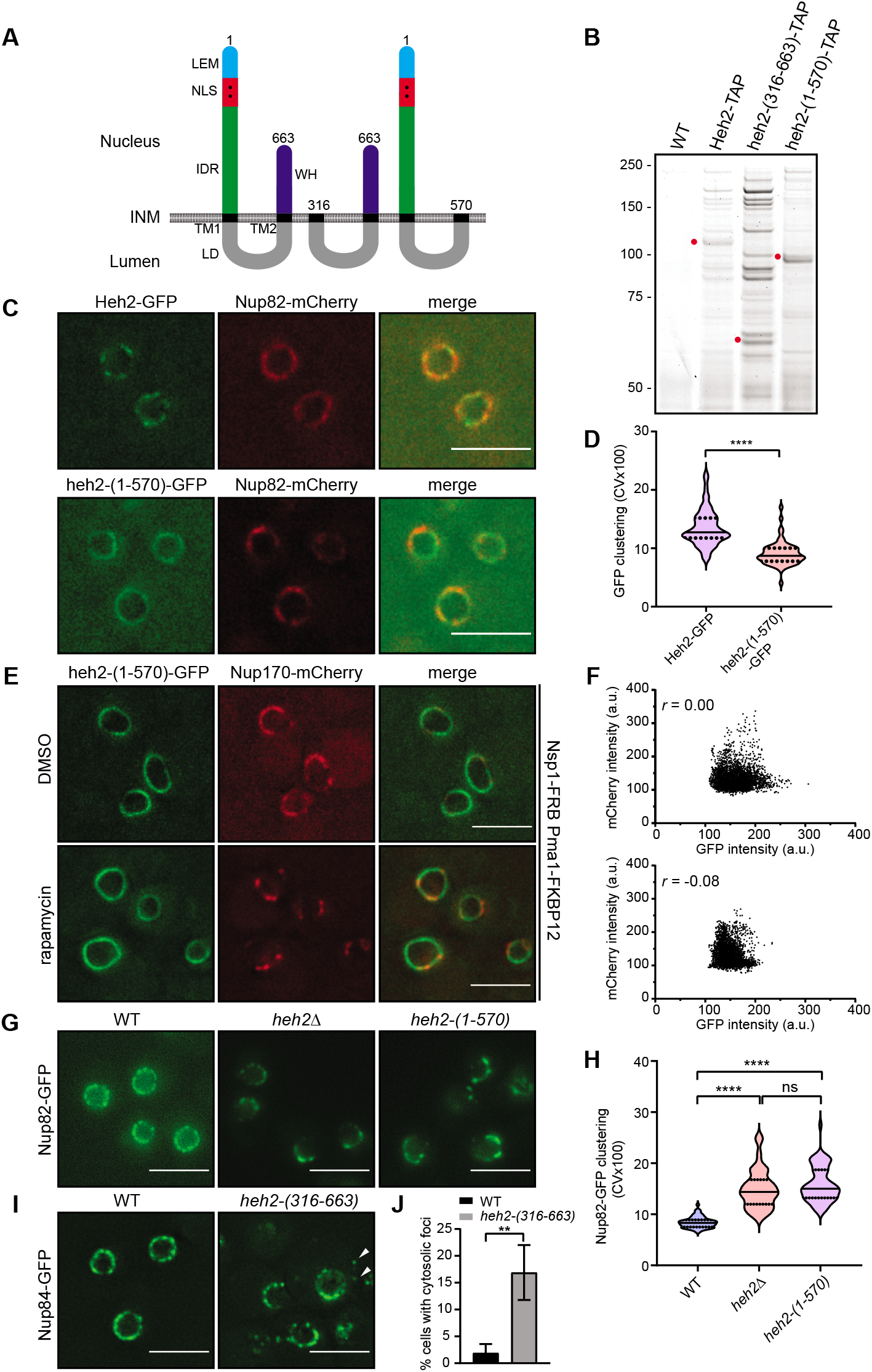
The WH domain of Heh2 is required for its association with NPCs. **(A**) Schematic of Heh2 and Heh2 truncations showing the LEM (Lap2-Emerin-Man1) domain, a bipartite nuclear localization signal (NLS), intrinsically disordered region (IDR), lumenal domain (LD), transmembrane domains (TM1 and TM2) and winged helix (WH); numbers represent amino acid numbers. INM, inner nuclear membrane. **(B)** Affinity purifications were performed from cell extracts derived from strains expressing the indicated TAP fusions or from WT cells (no TAP). Bound proteins were separated by SDS-PAGE and visualized by Coomassie staining. Numbers at left indicate position of MW standards in kD. Red circles denote position of TAP-fusions. **(C)** Deconvolved fluorescence micrographs of Heh2-GFP or heh2-(1-570)-GFP and the NPC marker Nup82-mCherry, with merge. Scale bar is 5 μm. **(D)** To quantitatively evaluate the distribution of Heh2-GFP and heh2-(1-570)-GFP, a coefficient of variation (CV) of the GFP fluorescence along the nuclear envelope was calculated. Individual CV values (multiplied by 100) were plotted with mean and SD from 60 cells, from three independent experiments. *p* values were calculated from Student’s t-test where **** indicates *p* ≤ 0.0001. **(E)** Deconvolved fluorescence micrographs of heh2-(1-570)-GFP and Nup170-mCherry with merge in cells expressing Nsp1-FRB and Pma1-FKBP12. Cells were treated with carrier (DMSO) or rapamycin. Addition of rapamycin leads to NPC clustering as described in Fig. 3A. Scale bar is 5 μm. **(F)** Scatterplot with Pearson correlation coefficient (*r*) of heh2-(1-570)-GFP and Nup170-mCherry fluorescence intensity (in arbitrary units, a.u.) along the nuclear envelope of 30 cells from three independent experiments like that shown in E. Values are from cells from DMSO (top) and rapamycin-treated (bottom) conditions. **(G)** The WH domain of Heh2 is required for normal NPC distribution. Deconvolved fluorescence micrographs of Nup82-GFP in indicated strain backgrounds. Scale bar is 5 μm. **(H)** To quantitatively evaluate the distribution of Nup82-GFP in the indicated strains, a coefficient of variation (CV) of the GFP fluorescence along the nuclear envelope was calculated. Individual CV values (multiplied by 100) were plotted with mean and SD from 60 cells, from three independent experiments. *p* values were calculated from one-way ANOVA with Tukey’s post-hoc test where ns is *p* > 0.05, \*\*\*\**p* ≤ 0.0001. **(I)** Deconvolved fluorescence micrographs of Nup84-GFP in WT and cells where *HEH2* is replaced by *heh2-(316-663)*. Arrowheads point to cytosolic Nup84-GFP foci. Scale bar is 5 μm. **(J)** Quantification of the percentage of cells where Nup84-GFP is found in the cytosol from experiment in I. Error bars are SD from four independent experiments. *p* values were calculated with unpaired t-test where ** indicates *p* ≤ 0.01.

### WH-domain-mediated interactions with NPCs are required for normal NPC distribution

As the Heh2 WH-domain was specifically required for Heh2-binding to NPCs, but not for INM targeting, there was an opportunity to define a putative NPC-specific function for Heh2. Indeed, deletion of *HEH2* leads to a marked clustering of Nup82-GFP, which was quantified as a coefficient of variation (CV) of the fluorescence along the nuclear envelope that was approximately double the value in WT cells (Fig. 6G, H). To directly test whether this phenotype was due to a loss of nup-binding, we assessed the distribution of Nup82-GFP in cells expressing *heh2-(1-570)*. Indeed, as shown in Fig. 6G, this targeted abrogation of the nup-binding WH domain also resulted in a clear redistribution of Nup82-GFP, showing a clustering coefficient nearly identical to that seen in *heh2*Δ cells (Fig. 6H). Thus, interactions between Heh2 and the NPC are required for normal NPC distribution.

Interestingly, expression of *heh2-(316-663)* from its endogenous locus also impacted NPC distribution, but with a unique phenotype. Because this truncation of Heh2 lacks its INM targeting information, this fusion will be mislocalized to the endoplasmic reticulum (King et al., 2006; Meinema et al., 2011). In these cells, Nup84-GFP accumulated in clusters at the nuclear envelope but also appeared within cytosolic foci (Fig. 6I, arrowheads) in ~17% of cells. Together then, these data support a model in which both the N-terminal and C-terminal domains of Heh2 are important for NPC distribution, however, the underlying mechanisms behind these alterations are unique and reflect either too little (in the case of heh2-(1-570)) and likely inappropriate (in the case of heh2-(316-663) interactions with nups.

## Discussion

We have explored the physical and functional relationship between the integral INM protein Heh2 and the NPC. This study was motivated by our prior discovery of predominantly genetic interactions between *HEH2* and nup genes (Yewdell et al., 2011), in addition to other work considering Heh2 as a factor in a NPC assembly surveillance pathway (Webster et al., 2014, 2016). In the latter, we imparted Heh2 the ability to discern between NPC assembly intermediates and fully formed NPCs. This concept was centered, in part, on data showing that Heh2 does not associate with clustered NPCs in *nup133*Δ strains, which was interpreted in a model where Heh2 does not bind to fully formed NPCs. We now provide a more nuanced explanation for these data, as deletion of Nup133 breaks Heh2’s otherwise robust physical association with the NPC (Fig. 5A). Thus, in light of the new data presented here, a reconsideration of the role of Heh2 in NPC biology is needed. Given these new observations, we suggest that Heh2 likely binds to fully formed NPCs. Several data support this assertion including: 1) The biochemical interactions that suggest the formation of a stable complex between Heh2 and the IRC (Fig. 1A, B). 2) The maintenance of these interactions even upon NPC assembly inhibition (Fig. 4C) and 3) The punctate distribution of Heh2 at steady-state and upon clustering of functional NPCs driven by the anchoring of Nsp1-FRB (Fig. 3C).

Despite the demonstration that Heh2 associates with NPCs, several new conundrums arise as a consequence of this work. The first is that we do not observe any robust physical association between Heh2 and the ORC, and yet, deletion of Nup133 leads to a loss of Heh2 binding to the NPC (Fig. 5A). In contrast, we cannot break Heh2’s association with NPCs by knocking out any individual component of the IRC (Fig. 5A, D). While the latter can be explained in a model where Heh2 makes several direct but redundant connections with nups, likely Pom152 and Nup170 and/or Nup188, the former is more challenging to interpret. Several potential models can be considered. The first deals with the very nature of *nup133*Δ NPC clustering, which has so far remained only partially explained on a mechanistic level. For example, one thought is that the association of NPCs with the pore membrane is destabilized without the amphipathic helix/ALPS motif in Nup133 (Drin et al., 2007), which may lead to pore clustering (Fernandez-Martinez et al., 2012). In such a scenario, given that it is an integral membrane protein, Heh2’s interactions with the NPC may depend on the presence of specific lipids or membrane curvature (or both) at the pore membrane. Alternatively, the clustering itself may sterically preclude an interaction with Heh2. It is also possible that the IRC may not be fully functional or be structurally perturbed in this context. Regardless of the underlying mechanism, as Heh2’s association with the NPC ultimately depends on the function of both of its major scaffold complexes (i.e. the IRC and ORC), we favor a model in which Heh2 can, through a mechanism that remains to be defined, “sense” the structural integrity of the NPC.

A model in which Heh2 is a sensor for the NPC scaffold fits within a quality control mechanism framework. For example, recent work suggests that NPC clustering can facilitate clearance of NPCs by autophagy (Lee et al., 2020). Thus, it is tempting to speculate that damage to the NPC scaffold may trigger the release of Heh2, which would in turn lead to the clustering of damaged NPCs. Such an idea is supported by the clustering that we observe in contexts where Heh2-NPC interactions are abrogated (Fig. 6G, H). Similarly, as we have previously reported, NPC clustering may also be an input that ensures that damaged or malformed NPCs are not transmitted to daughter cells (Webster et al., 2014). Thus, the consistent theme is that breaking interactions between Heh2 and NPCs may be an input to their segregation and/or clearance. A corollary to this is that Heh2 bound to NPCs may in fact promote the inheritance of functional NPCs. This may be best illustrated by work from *S. japonicus* where it was demonstrated that the Heh2 orthologue contributes to anchoring NPCs to chromatin to promote their proper segregation between daughters (Yam et al., 2013). Indeed, our observation that Heh2 also engaged in interactions with the IRC in *S. pombe* argues that it supports a fundamental role(s) across diverse yeasts.

How, then, do interactions between Heh2 and NPCs ensure proper NPC distribution? We speculate that in the absence of mechanisms to keep NPCs apart, NPCs have an inherent conformation or affinity that drives their clustering. In this scenario, binding NPCs to INM proteins could help ensure their physical segregation. Although this could be envisaged purely as a steric inhibition of NPC-NPC interactions, we favor the concept that the distribution of NPCs and other elements of the nuclear architecture are co-dependent. Indeed, our prior work suggests that SpHeh2 antagonizes the flow of chromatin into nuclear deformations (Schreiner et al., 2015), in essence maintaining normal chromatin distribution at the nuclear periphery, a direct corollary of the effect here on NPC distribution. As SpHeh2 binds both chromatin (Gonzalez et al., 2012; Steglich et al., 2012) and NPCs (this work), it is tempting to speculate that it supports the normal organization of NPCs and chromatin by dynamically linking these two major structural components of the nucleus. This concept is consistent with evidence in mammalian cells where NPCs are well established to be anchored to the lamin network (Daigle et al., 2001; Maeshima et al., 2006; Xie and Burke, 2017; Kittisopikul et al., 2020). In scenarios in which this lamin connection is broken, for example in lamin knockouts, NPCs also cluster together (Xie and Burke, 2017; Kittisopikul et al., 2020). Although NPCs are more dynamic along the nuclear envelope in budding yeast (Belgareh and Doye, 1997; Bucci and Wente, 1997), their interactions with chromatin through multiple mechanisms (Luthra et al., 2007; Tan-Wong et al., 2009) could nonetheless contribute to their normal distribution. Whether clustering has an impact on NPC function per se remains ill defined, although one could speculate that NPC clustering has a more profound impact on the NPC’s roles in chromatin organization and gene expression as opposed to nuclear transport (Capelson et al., 2010; Raices and D’Angelo, 2017).

One particularly interesting feature of our analysis of Heh2 is that the NPC binding and INM targeting sequences are distinct and on two physically separated domains. Certainly there is evidence from both genetic and biochemical analyses where the function of specific domains of the LEM domain proteins can be separated (Grund et al., 2008; Yewdell et al., 2011; Barrales et al., 2016; Hirano et al., 2018; Thaller et al., 2019; von Appen et al., 2020). However, we wonder whether there are functional implications for the integration of these two interaction platforms, which could place Heh2 in a tug-of-war between its residence bound to the NPC and its release to the INM. This would be yet another example in an emerging theme for these LEM domain proteins in which they bridge distinct sets of physical interactions to maintain the dynamic organization of the nuclear envelope system.

## Materials and methods

### Yeast culture and strain generation

All yeast strains used in this study are listed in Table S1. *S. cerevisiae* strains were grown in YPD consisting of 1% Yeast extract (BD), 2% Bacto-peptone (BD) and, 2% D-glucose (Sigma). For microscopy experiments, YPD was supplemented with 0.025% adenine hemi-sulfate (Sigma). Yeast cells were grown at 30°C to mid-log phase, unless otherwise stated. Transformation of *S. cerevisiae* cells, mating, sporulation and tetrad-dissections were carried out using standard protocols (Amberg et al., 2005). Deletion and truncation of yeast ORFs and tagging of ORFs with fluorescent protein genes, FRB and TAP-tags was performed utilizing the pFA6a or pK3F plasmid templates (Longtine et al., 1998; Zhang et al., 2017).

*S. pombe* strains were grown in YE5S media consisting of 5% Yeast extract (BD), 30% D-glucose (Sigma) and 1.25% SP complete supplements (adenine hemisulfate, L-histidine hydrochloride monohydrate, L-leucine, L-lysine hydrochloride and uracil) from Sunrise Science products, at 30°C. *S. pombe* strains were crossed and maintained utilizing standard media and techniques as described in (Moreno et al., 1991). PCR based gene disruption and tagging were performed utilizing pFA6a plasmid templates (Bähler et al., 1998; Hentges et al., 2005).

### Plasmids

All plasmids used in this study are listed in Table S2. The pFA6a-TAP-his3MX6 and pFA6a-TAP-TRP1 plasmids were constructed as follows: the TAP coding sequence was PCR-amplified from chromosomal DNA from a strain expressing Heh2-TAP (SBCPL42, Dharmacon yeast resources) using Phusion High fidelity DNA polymerase (New England BioLabs) and cloned into the *PacI* and *AscI* sites of pFA6a-his3MX6 and pFA6a-TRP1.

pFA6a-3xHA-FRB-GFP-his3MX6 was generated by Gibson Assembly (New England BioLabs). The 3xHA epitope coding sequence was PCR-amplified from pFA6a-3xHA-hisMX6 (Longtine et al., 1998) using Q5 DNA polymerase (New England BioLabs) and assembled into pFA6a-FRB-GFP-hisMX6, or pFA6a-FRB-hisMX6 (EUROSCARF) digested with *SalI* and *PacI*.

### Immunoaffinity purification

To affinity purify TAP-fusions, *S. cerevisiae* strains were grown overnight and 2 ml of culture was diluted into 1 l of YPD the next morning and grown for 20-24 h to late log phase (OD600 ~2). *S. pombe* cells were grown overnight and transferred to fresh medium the next morning to an OD600 of 0.1 and grown for 7 h. *S. pombe* cells were further diluted to an OD600 of 0.01 in 1 l YES medium and grown for another 18-20 h. Both *S. cerevisiae* and *S. pombe* cells were grown at 30°C at 200 rpm and cells were harvested by centrifugation. Cells were washed with ice-cold water once, collected by centrifugation and resuspended in 100 μl freezing solution (20 mM HEPES, pH 7.4, 1.2% polyvinylpyrrolidone and protease inhibitor cocktail [Sigma]) per g of cells. The cell slurry was snap-frozen in liquid nitrogen immediately. The frozen cell pellets were cryo-milled 6 times at 30 Hz for 3 min in a Retsch MM400 mixer mill and stored at −80°C.

To perform immunoaffinity purifications, 200 mg of frozen yeast grindate was resuspended in 4-times volume of homogenization buffer (400 mM Na3Cit, pH 8.0, 10 mM Deoxy Big CHAP) and protease inhibitor cocktail at room temperature. The homogenate was clarified by centrifugation at 16,000 g for 10 min at 4°C. The soluble fraction was incubated with 25 μl of Rabbit-IgG coated Dynabeads for 1 h at 4°C under gentle rotation. After binding, beads were collected on a magnetic rack and washed three times with 500 μl ice-cold homogenization buffer. The proteins were eluted by incubating beads with 20 μl of 1X NuPAGE lithium dodecyl sulfate sample buffer (Invitrogen) at room temperature for 10 min. The eluate was separated on a magnetic rack and further incubated with 50 mM DTT at 70°C for 10 min. The eluted proteins were separated on a 4-12% NuPAGE gel (Novex) and stained with Imperial protein stain (Thermo Scientific). The proteins of interest were excised for identification by MS.

### Conjugation of Dynabeads with Rabbit IgG

Purified rabbit IgG (Sigma, I5006) was dissolved in 0.1 M sodium phosphate buffer, pH 7.4, to a final concentration of 1 mg/ml. The IgG solution was filtered through a 0.22 μm syringe filter and mixed with an equal volume of 3 M (NH4)2SO4. For conjugation, 100 mg of Dynabeads^®^ M-270 Epoxy (Invitrogen) were transferred to a 15 ml centrifuge tube, suspended in 6 ml 0.1 M sodium phosphate buffer, pH 7.4 and incubated at room temperature for 15 min on a tube rotator. The beads were collected on a magnetic rack, the buffer aspirated and beads were washed again with 0.1 M sodium phosphate buffer, pH 7.4 by vortexing. The buffer was removed and beads were resuspended in 2 ml of IgG solution and incubated at 30°C for 65-70 h on a tube rotator. The beads were separated on a magnetic rack and quickly washed with 100 mM glycine, pH 2.5, followed by a wash with 10 mM Tric-HCl, pH 8.8. Beads were again washed quickly with freshly prepared 100 mM Triethylamine and followed by 4 washes with PBS for 5 min each and one wash with PBS with 0.5% Triton X-100 for 15 min. The beads were washed one final time with PBS, collected on a magnetic rack and resuspended in 667μl PBS with 50% glycerol.

### Anchor-away experiments

The anchor-away experiments were performed as described by Haruki et al., 2008. Briefly, strains expressing Nup-FRB fusions and Pma1-FKPB12 in HHY110 (*tor1-1 fpr1Δ*) were incubated with a final concentration of 1 μg/ml rapamycin for 30 min (to cluster NPCs in the context of Nsp1-FRB) or 3 h to inhibit assembly (Nup192-FRB).

### Fluorescence microscopy, image processing and analysis

Fluorescence micrographs were acquired on a DeltaVision microsope (Applied Precision, GE Healthcare) with a 100x, 1.4 NA objective (Olympus). The images were captured with a CoolSnapHQ^2^ CCD camera (Photometrics). Fluorescence micrographs were deconvolved with the iterative algorithm sofWoRx. 6.5.1 (Applied Precision, GE Healthcare).

Clustering of NPCs was quantified as described previously (FernandezMartinez et al., 2012): A 6-pixel wide freehand line was drawn along the nuclear envelope contour and mean fluorescence intensities were measured using FIJI/ImageJ (Schindelin et al., 2012). Clustering was assessed by calculating the coefficient of variance (SD/mean X 100) of the fluorescence intensities at the nuclear envelope.

### Modeling of NPC spokes

Color coding of an isosurface representation of individual nup densities as assigned in Kim et al. 2018 within an individual spoke of the NPC from the PDB DEV ID:00000010 was completed using ChimeraX (UCSF) (Goddard et al., 2018).

## Acknowledgements

We thank Jean Kanyo and the Yale Keck Biotechnology Resource Laboratory for help with MS analysis. We would like to thank for Valérie Doye for *S. pombe* strains. We would like to thank the members of the Lusk and King laboratories for critical input on experimental design and data analysis. This work was supported by the NIH: R01 GM105672 to CPL, R21 HG006742 to CPL and MCK, and R01 GM112108 and P41 GM109824 to MPR and the NSF: EFMA-1806504 to MCK.

---

**Table S1.**
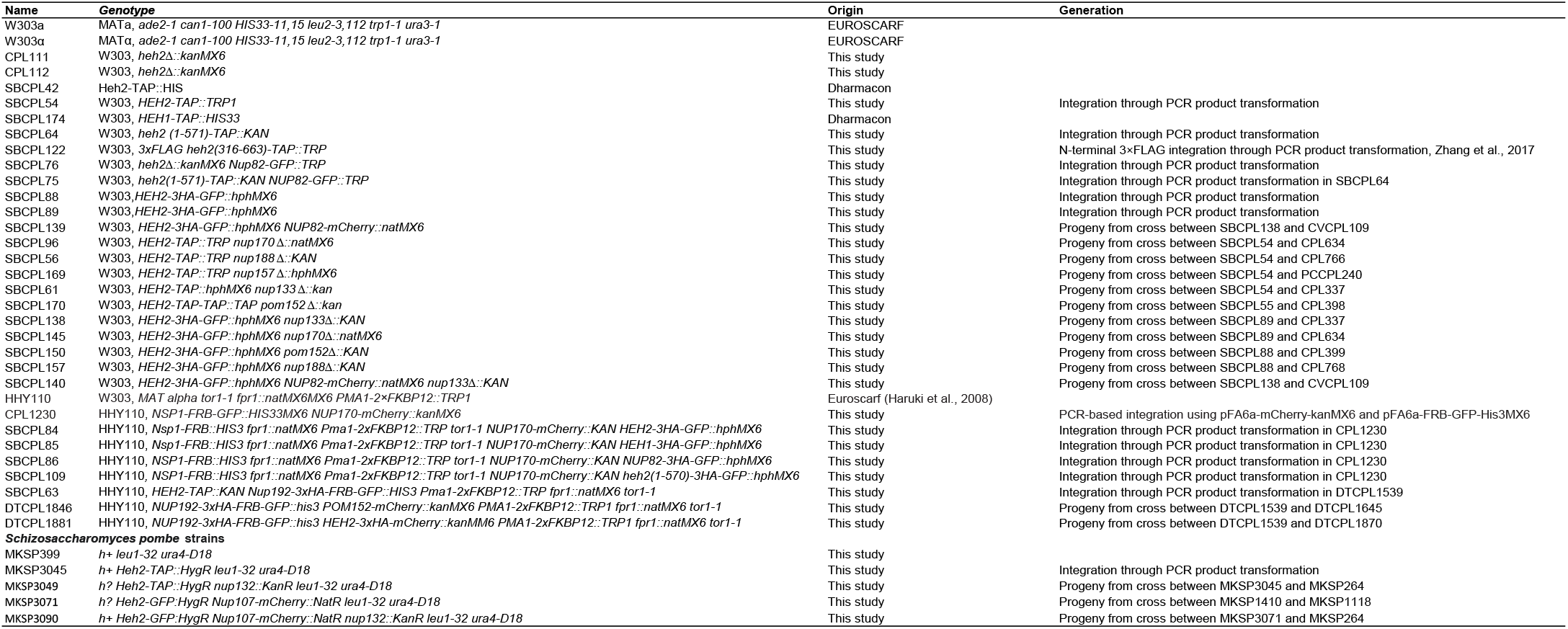
Yeast strains *Saccharomyces cerevisiae* strains

**Table S2.** Plasmids

| Name | Description | Source |
| --- | --- | --- |
| pFA6a-GFP-his3MX6 | Template for PCR based chromosomal integration of GFP ORF | Longtine et al., 1998 |
| pFA6a-GFP-natMX6 | Template for PCR based chromosomal integration of GFP ORF | Van Driessche et al., 2005 |
| pFA6a-GFP-kanMX6 | Template for PCR based chromosomal integration of GFP ORF | Longtine et al., 1998 |
| pFA6a-hphMX6 | Template for PCR based chromosomal integration of hphMX6 cassette | Longtine et al., 1998 |
| pFA6a-natMX6 | Template for PCR based chromosomal integration of natMX6 cassette | Longtine et al., 1998 |
| pFA6a-kanMX6 | Template for PCR based chromosomal integration of kanMX6 cassette | Longtine et al., 1998 |
| pFA6a-mCherry-kanMX6 | Template for PCR based chromosomal integration of mCherry ORF | EUROSCARF |
| pFA6a-mCherry-natMX6 | Template for PCR based chromosomal integration of mCherry ORF | EUROSCARF |
| pSBCPL3 | pFA6a-TAP-his3MX6, template for PCR based chromosomal integration of TAP-TAG | This study |
| pSBCPL4 | pFA6a-TAP-TRP, template for PCR based chromosomal integration of TAP-TAG | This study |
| pK3F | N-ICE plasmid pK3F, for N-terminal 3×FLAG integration | Addgene |
| pSH47 | Cre recombinase under the *GAL1* promoter | EUROSCARF |

## Notes

### Competing Interest Statement

The authors have declared no competing interest.

